# Evolution of genetic variance during adaptive radiation

**DOI:** 10.1101/097642

**Authors:** Greg M. Walter, J. David Aguirre, Mark W. Blows, Daniel Ortiz-Barrientos

**Affiliations:** University of Queensland, School of Biological Sciences, St. Lucia QLD 4072, Australia; Massey University, Institute of Natural and Mathematical Sciences, Auckland 0745, New Zealand

**Keywords:** adaptive radiation, genetic constraint, additive genetic variance, phenotypic, divergence, covariance tensor

## Abstract

Genetic correlations between traits can bias adaptation away from optimal phenotypes and constrain the rate of evolution. If genetic correlations between traits limit adaptation to contrasting environments, rapid adaptive divergence across a heterogeneous landscape may be difficult. However, if genetic variance can evolve and align with the direction of natural selection, then abundant allelic variation can promote rapid divergence during adaptive radiation. Here, we explored adaptive divergence among ecotypes of an Australian native wildflower by quantifying divergence in multivariate phenotypes of populations that occupy four contrasting environments. We investigated differences in multivariate genetic variance underlying morphological traits and examined the alignment between divergence in phenotype and divergence in genetic variance. We found that divergence in mean multivariate phenotype has occurred along two major axes represented by different combinations of plant architecture and leaf traits. Ecotypes also showed divergence in the level of genetic variance in individual traits, and the multivariate distribution of genetic variance among traits. Divergence in multivariate phenotypic mean aligned with divergence in genetic variance, with most of the divergence in phenotype among ecotypes associated with a change in trait combinations that had substantial levels of genetic variance in each ecotype. Overall, our results suggest that divergent natural selection acting on high levels of standing genetic variation might fuel ecotypic differentiation during the early stages of adaptive radiation.

## Introduction

Evolutionary biologists have long sought to understand the processes that have created the dramatic diversification of species we see in nature (1). Adaptive radiation is one process that drives the creation of biological diversity and occurs when groups of organisms colonize and rapidly adapt to heterogeneous environments, leading to divergence and speciation (2). Differences in directional selection between environments can favour adaptive phenotypic divergence between populations and lead to the formation of ecotypes (3, 4), provided sufficient genetic variation is present in populations exposed to spatial variation in natural selection (5). However, experimental work on understanding how spatial variation in natural selection creates adaptive radiation across a heterogeneous landscape is rare (e.g., 6, 7, 8).

Replicate populations that occupy similar environments and possess similar phenotypes can be used as natural experiments to explore how genetic and phenotypic variation has evolved during adaptive divergence. Populations that experience similar environments often evolve similar phenotypes, creating strong correlations between habitat and morphology (9–11). Populations adapted to similar environments may evolve specific combinations of traits, creating discontinuous phenotypes between contrasting environments across a heterogeneous landscape (12). For example, stickleback fish have exhibited rapid morphological adaptive divergence into a novel environment via the repeated fixation of alleles from standing genetic variation (13, 14), and *Anolis* lizards have repeatedly evolved specialized limbs in similar habitats on different islands (7, 15, 16).

Correlations between habitat and morphology are often examined on single traits in isolation, but natural selection typically acts on multiple traits simultaneously (17–20). Consequently, adaptation can favour the evolution of beneficial combinations of traits within an environment and create adaptive divergence in multivariate phenotypes between contrasting environments (21–23), potentially underlying the origin of ecotypes during adaptive radiation (2, 3, 24). Since populations exposed to the same environment can also diverge as a consequence of random drift, quantifying ecotypic divergence requires comparing the divergence in multivariate phenotypes among ecotypes to differences among replicate populations within an ecotype (25–27). Stronger morphological divergence between ecotypes from contrasting environments than between populations within environments suggests that differences in the environment have promoted divergence (16, 23, 28), and such experimental systems can be used to identify how divergent natural selection has created adaptive morphological diversification during an adaptive radiation (29).

The magnitude of additive genetic variance in the direction of selection determines the rate of adaptive phenotypic evolution, but the availability of genetic variance in multivariate phenotypes depends on the extent of genetic correlation between traits (25,27, 30–32). Genetic correlations among traits concentrate genetic variation into particular trait combinations at the expense of other trait combinations (33). When genetic correlations bias the distribution of genetic variation away from the direction of selection, constraints on the rate of adaptive evolution are likely (34). The additive genetic variance-covariance matrix (**G**) summarises the genetic relationships between traits, providing the framework to investigate multivariate phenotypic evolution (27).

Adaptation is expected to bias evolution along the multivariate axis of greatest genetic variance (***g***_max_) (2, 28, 29, 35) at least in the early stages of divergence as there is little reason to expect that the major axis of additive genetic variance will be in a similar direction to that of natural selection following colonization of a novel environment (29).

In practice, not only will divergent selection change the phenotypic mean, but it may also result in the evolution of the genetic variance underlying traits during adaptation. For example, rare alleles held at mutation selection balance may become beneficial when exposed to a new environment, rapidly altering the phenotype mean as well as the distribution of genetic variance (36, 37). If genetic variance can align with the direction of natural selection, then adaptation from standing genetic variation may explain how rapid divergence occurs during adaptive radiation. In attempting to characterize an adaptive radiation, ideally we would understand what divergence in multivariate mean phenotype has occurred, explore how genetic variance underlying multivariate phenotypes has evolved and then quantify how changes in genetic variance align with changes in multivariate phenotypic mean. Here, we report a set of experiments that enabled us to determine how the phenotypic mean and genetic variance have diverged in the early stages of an adaptive radiation.

We investigated the adaptive radiation of *Senecio lautus*, an herbaceous wildflower across a heterogeneous landscape. *Senecio lautus* is a species complex native to Australia, New Zealand and several Pacific Islands. Although *S. lautus* contains many taxonomic species, we focussed on varieties within *S. pinnatifolius*, which are native to Australia and occupies a diverse array of habitats. We investigate divergence of populations from four distinct habitats; coastal headlands (*S. pinnatifolius* var. *maritimus*; Headland ecotype), coastal sand dunes (*S. pinnatifolius* var. *pinnatifolius*; Dune ecotype), moist tableland rainforests (*S. pinnatifolius* var. *serratus*; Tableland ecotype) and dry sclerophyll woodland (*S. pinnatifolius* var. *dissectifolius*; Woodland ecotype) (38, 39). Ecotypes from these habitats display strong morphological differentiation associated with the different environments and plants maintained their field morphology when grown under common garden conditions, indicating that phenotypic differences between populations have a strong underlying genetic basis (38–41). Transplant experiments have revealed that these ecotypes are adapted to their local environments (42–44), where extrinsic reproductive isolation is strong, but intrinsic reproductive isolation is weaker, suggesting barriers are largely geographic and ecological (42–45). Phylogenetic and population genetic analyses have revealed two major clades of taxa of low genetic differentiation yet high morphological diversity. Divergence between clades is less than one million years, and some ecotypes (coastal and alpine) have formed independently multiple times (46, 47), suggesting that, similar to other systems (e.g., African cichlids (48) and the Caribbean *Anolis* lizards (16)) the *S. lautus* species complex has recently undergone parallel evolution and adaptive radiation.

To investigate how divergence in mean multivariate phenotype has occurred within and between ecotypes, we sampled seeds from replicate populations of each of the four ecotypes, which were then grown under common garden conditions. In these populations we measured ten traits related to plant architecture and leaf morphology. Using a breeding design, we then estimated additive genetic (co)variance matrices for each ecotype for the same ten morphological traits and quantified divergence in genetic variance between ecotypes. Finally, we investigated whether divergence in genetic variance and divergence in phenotypic mean aligned to investigate how adaptive divergence has occurred during adaptive radiation.

## Results

### Glasshouse experiments

We collected seeds from four populations for each ecotype (Supplementary Table S1). Populations occupy small patches of habitat, which restricted us to sampling seeds from 30-45 individuals per population. Seeds were taken from individuals at ten metre intervals to reduce the risk of sampling close relatives. To compare divergence in phenotype mean with divergence in genetic variance we conducted two separate glasshouse experiments. In the first experiment we grew 16 individuals from four populations of each ecotype (ecotype n = 4; population n = 16; total n = 242), which we used to estimate divergence in multivariate phenotype mean. In the second experiment we used a North Carolina II breeding design to estimate genetic variance components for two populations of each ecotype (ecotype n = 4; population n = 8; total individuals = 1,259).

To estimate genetic variance we grew two generations of plants in the glasshouse. To establish the parent generation we grew seeds collected from the natural populations and crossed them using a breeding design. Only one seed from each individual sampled in the field was grown because seeds taken from the same plant were likely pollinated by different individuals, making parentage uncertain. Within each population half the individuals were designated sires (n = 17-23 per population; Supplementary Table S2) and the other half dams (n = 15-23 per population; Supplementary Table S2). Each sire was then randomly crossed to two dams according to a North Carolina II breeding design, where variance between paternal half-siblings represent one quarter of the additive genetic variance (49). Three to four offspring for each full-sibling family produced from these crosses were grown and phenotyped in a glasshouse experiment conducted in 2015, which totalled 934 individuals.

Estimating genetic variance requires large sample sizes and breeding designs, which logistically restricted us to only two populations for each ecotype. However, measuring divergence in multivariate phenotype mean is best achieved using a hierarchical framework by assessing differences between ecotypes, taking into account differences between populations within ecotypes. We used the first experiment to represent divergence in mean multivariate phenotype because it contained four populations, providing three degrees of freedom, while the second experiment only provided one degree of freedom. However, differences in mean phenotypes between populations grown in both experiments did not change considerably (Supplementary Figure S3), verifying the use of the first glasshouse experiment to represent divergence in phenotype mean for the second experiment.

### Divergence in multivariate mean phenotype

The four ecotypes displayed visually striking differences in leaf morphology (Figure 1A) and plant architecture (Figure 1B) under common garden conditions, suggesting a strong association between morphology and habitat. A multivariate analysis of variance (MANOVA) indicated there was significantly more variation between ecotypes than could be accounted for by variation within ecotypes (Wilks’ lambda λ = 1.09x10^-04^, *F*_3_,_12_ = 6.76, *P* = 0.002). Multivariate phenotypic divergence between ecotypes explained much more of the total variance (62%) than divergence between populations within ecotypes (11%), highlighting a strong and consistent pattern of ecotypic divergence. From the MANOVA we calculated the phenotypic divergence matrix (**D** matrix) of population means, which described divergence between ecotypes in multivariate space (27, 28, 50). The first eigenvector of **D** (***d***_max_) explained 67% of the divergence in multivariate mean phenotype, created by differences in traits relating to plant size in one direction and leaf shape in the other direction (Figure 1C), and separated the Tableland and Woodland ecotypes (Figure 1D). The second vector (***d***_2_) explained 32% of the variation, created by plant size and leaf complexity in one direction, and number of indents and circularity in the other direction (Figure 1C), and separated the Headland and Woodland from the other ecotypes (Figure 1D). The third eigenvector (***d***_3_) explained only 3% of divergence (Figure 1C), and was primarily associated with the number of branches.

**Figure 1.**
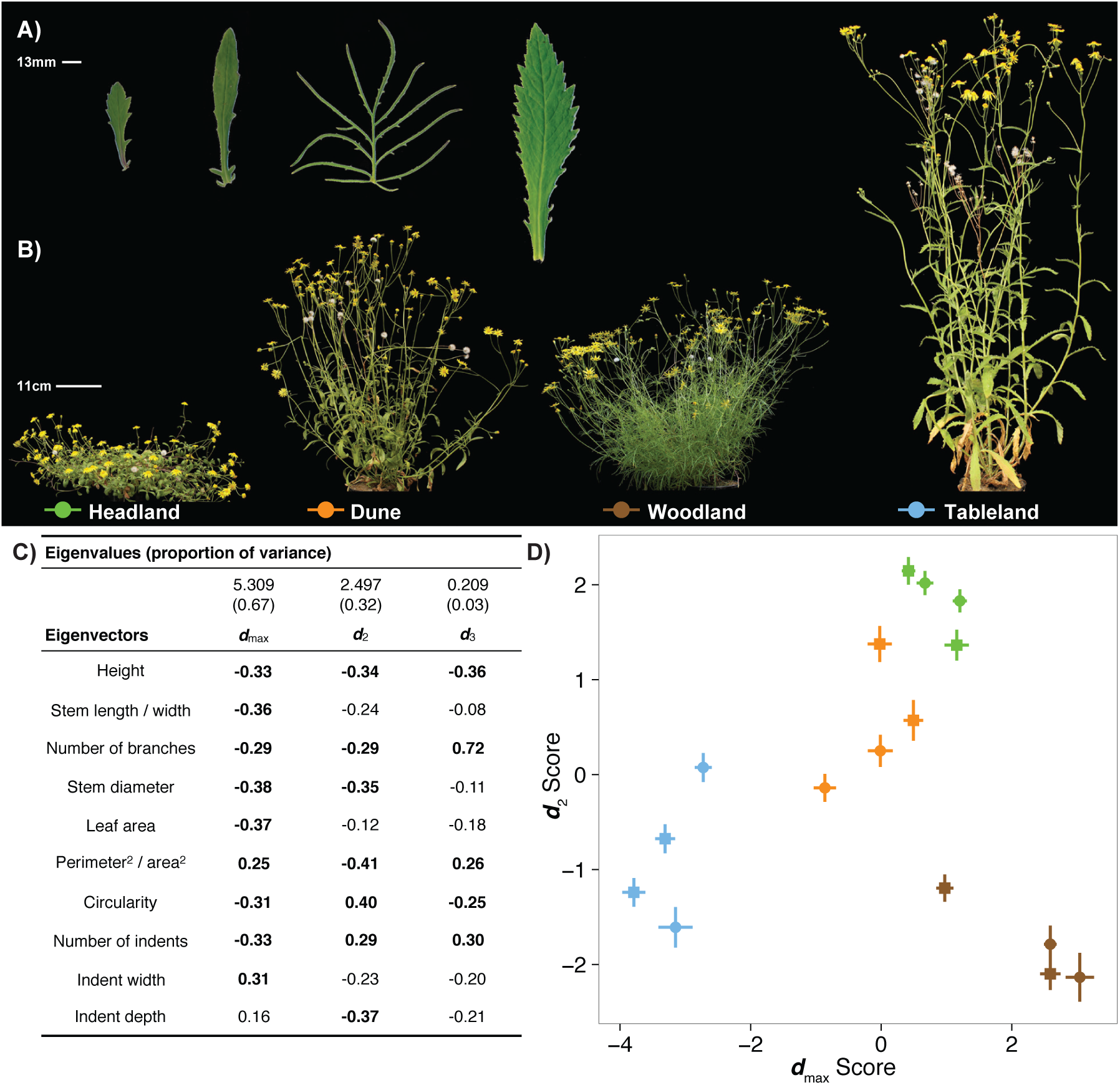
Ecotypes show strong differences in **(A)** leaf morphology and **(B)** plant architecture. **(C)** Eigenanalysis of **D** showed that multivariate divergence in mean occurred in two major axes created by different combinations of plant architecture and leaf traits. Numbers in bold represent trait loadings higher than 0.25. **(D)** The score for each population calculated from the eigenanalysis of **D** showed that populations from the same ecotype group together. The first eigenvector separated Tableland and Woodland from the remaining ecotypes, while the second eigenvector separated the Woodland and Headland from the remaining ecotypes.

### Genetic variance underlying plant morphology

Many of the observed heritabilities for univariate traits exceeded the magnitude of sampling error in our experimental design, although there was considerable variation in the magnitude of heritability among populations (Figure 2). For example, the Headland ecotype displayed high (h^2^ > 0.4) and significant heritabilities for architecture traits and leaf area, while the same traits in the Woodland ecotype had lower heritabilities that did not exceed sampling error in 4 of 5 cases (Figure 2).

**Figure 2.**
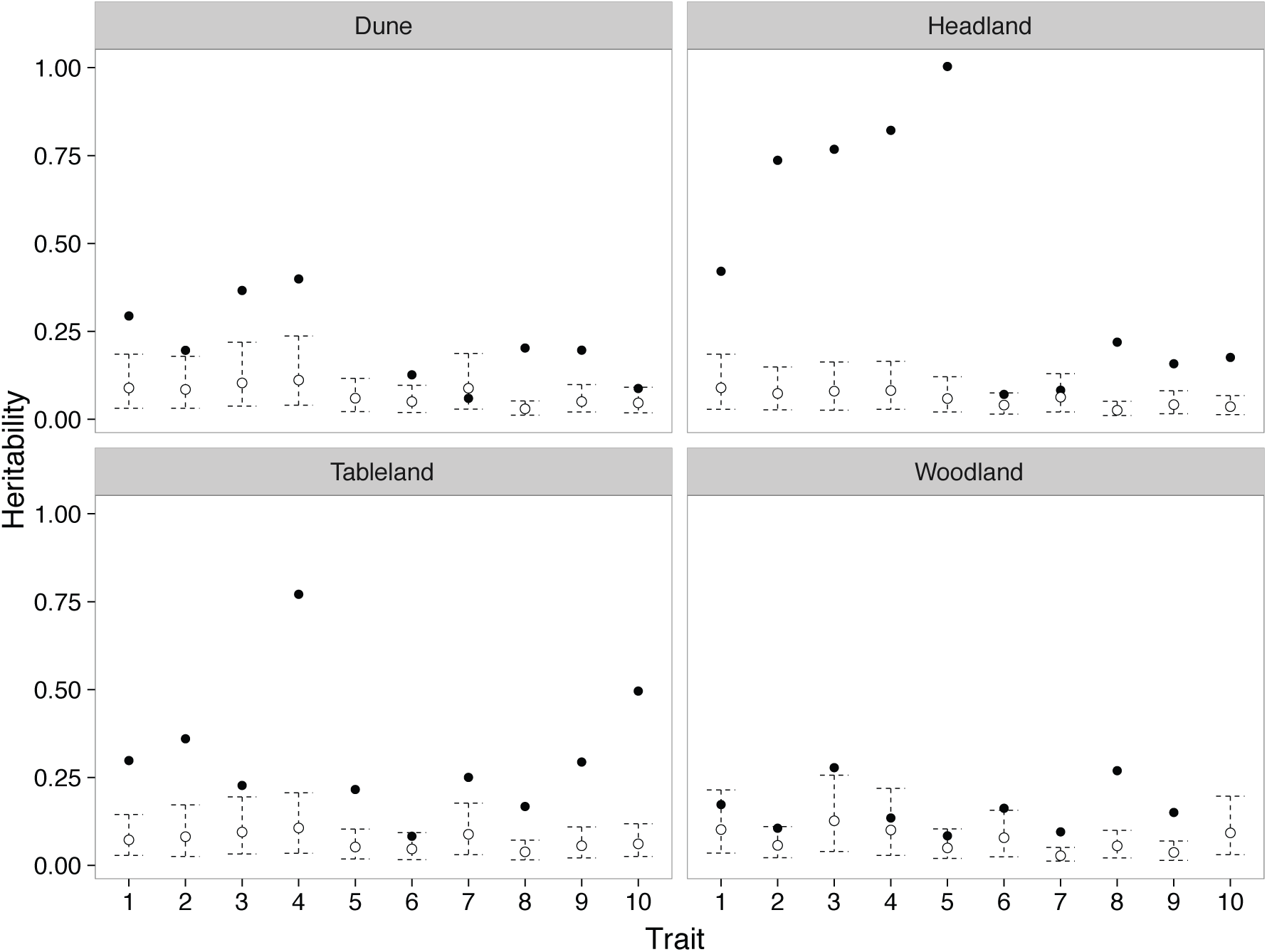
Observed heritabilities (filled circles) compared to the distribution of randomly estimated heritabilities (unfilled circles). Dashed lines represent 95% HPD intervals. Architecture traits showed very high heritabilities in the Headland and Tableland, while heritabilities for all traits were much lower in the Dune and Woodland ecotypes. Traits: 1 = Height, 2 = Stem length / width, 3 = # branches, 4 = Stem diameter, 5 = Leaf area, 6 = Perimeter^2^ / Area^2^, 7 = Circularity, 8 = # indents, 9 = Indent width, 10 = Indent depth.

Genetic correlations tended to be positive among architecture traits, but negative among leaf shape traits (Supplementary Table S4). Genetic correlations between these two trait types tended to be negative. Overall, the magnitude of genetic correlations were relatively weak, which was reflected in the relatively low proportion of genetic variance accounted for by the leading eigenvalue of G in each ecotype (Table 1). The Headland ecotype was the only ecotype that had > 50% of the genetic variance in ***g***_max_, while the other ecotypes showed a much more uniform distribution of genetic variance across eigenvectors (Table 1 and Figure 3). Visual inspection of the eigenvectors (Table 1) suggested a consistent pattern in three of the ecotypes, where the linear combinations of ***g***_max_ had higher loading from architectural traits, while linear combinations with stronger contributions from leaf traits captured smaller amounts of genetic variance (Table 1). The Woodland ecotype was the exception, where both architectural and leaf traits were represented in eigenvectors containing both high and low genetic variance. Most eigenvalues of **G** were significantly higher than expected from sampling error in all four ecotypes (Figure 3).

**Table 1.**
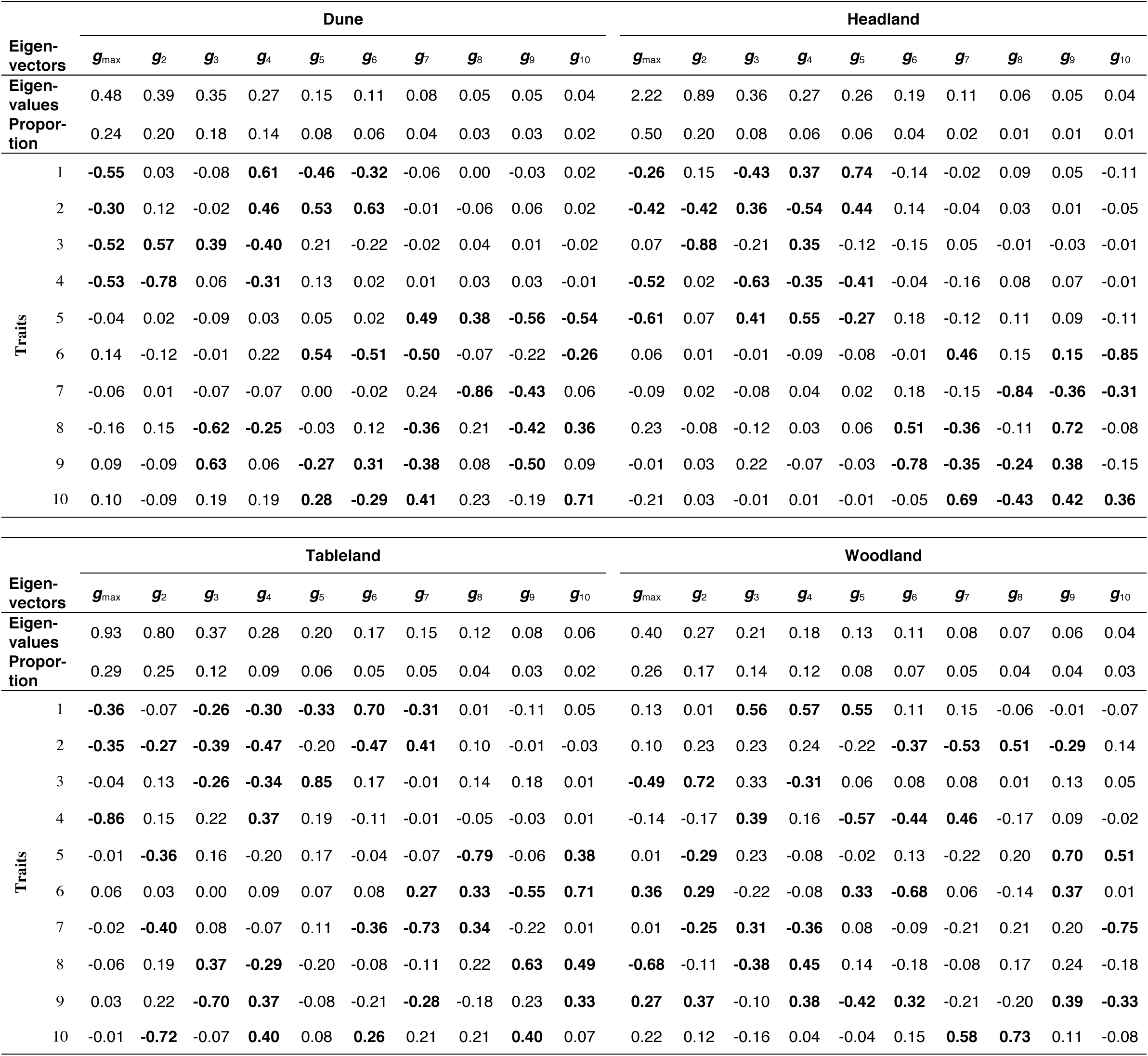
Eigenanalysis of **G** showed that architecture traits contributed to the genetic variance contained in ***g***_max_ for all ecotypes except the Woodland. Leaf shape traits were only represented in eigenvectors with low genetic variance. Numbers in bold denote trait loadings higher than 0.25. Traits: 1 = Height, 2 = Stem length / width, 3 = # branches, 4 = Stem diameter, 5 = Leaf area, 6 = Perimeter^2^ / Area^2^, 7 = Circularity, 8 = # indents, 9 = Indent width, 10 = Indent depth.

**Figure 3.**
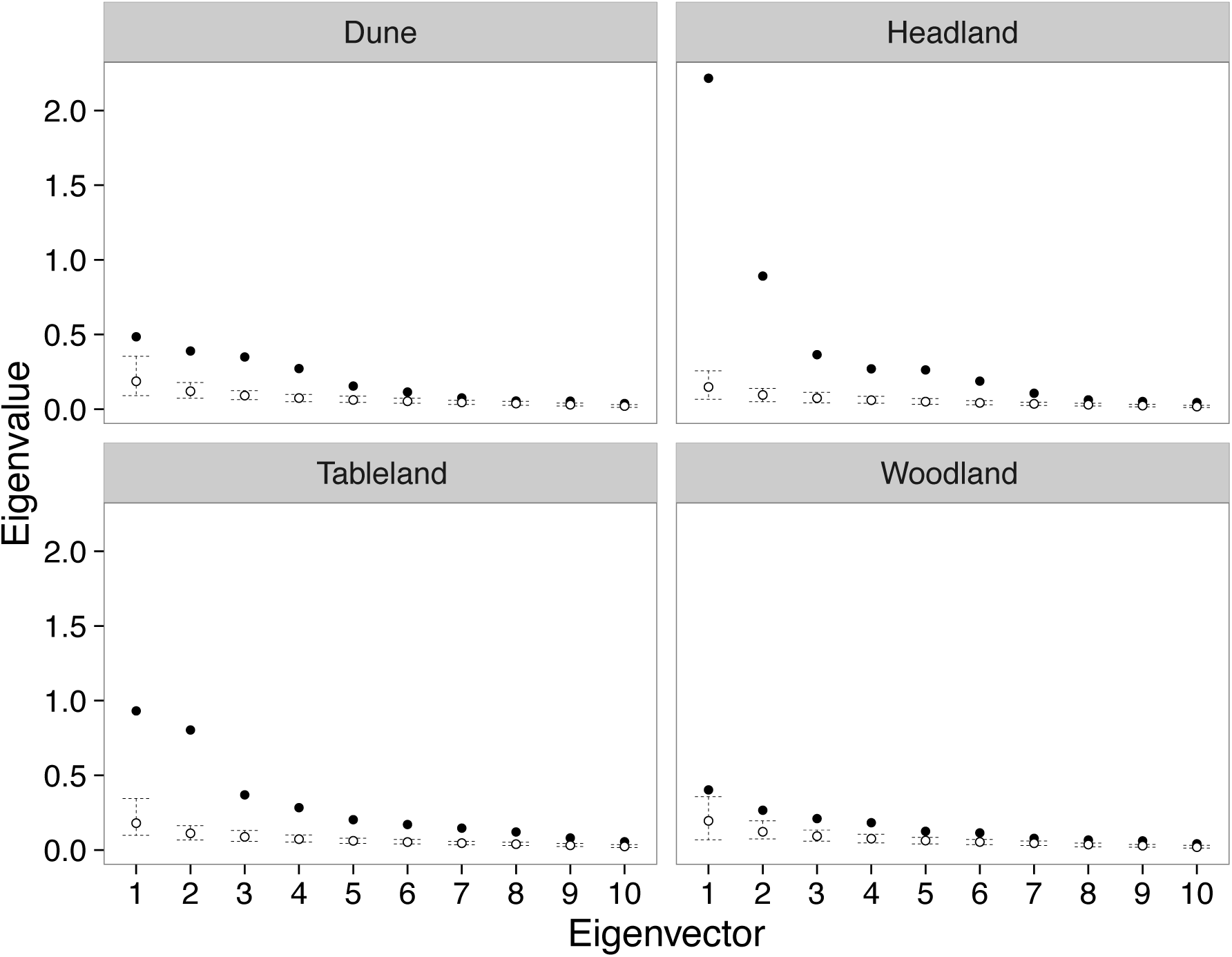
Observed eigenvalues of **G** (filled circles) compared to the distribution of random eigenvalues of **G** (unfilled circles). Dashed lines represent 95% HPD intervals. The Headland was the only ecotype that showed a much higher amount of variance in ***g***_max_, while the remaining ecotypes exhibited a similar partitioning of genetic variance among eigenvectors.

### Divergence in multivariate genetic variance among ecotypes

To compare differences in **G** among ecotypes we used a genetic covariance tensor. We found three significant eigentensors of **G** that explained 38%, 10% and 7% of the divergence in genetic variance between ecotypes, respectively. All three eigentensors described more divergence in genetic variance than expected by sampling error alone (Supplementary Figure S5). The coordinates of the ecotypes in the space of the eigentensor revealed how each ecotype contributed to differences in genetic variance. We were only able to detect significant differences between ecotypes in the first eigentensor (E1) (Supplementary Figure S6). We therefore concentrate our analysis on the first eigentensor (Table 2), while the full tensor analysis can be found in Supplementary Table S7

**Table 2.** Table summary of the covariance tensor analysis of **G**, α denotes the magnitude of the difference between matrices described by the eigentensor, with the proportion of variance in parentheses. Eigenvector and λ represents the eigenanalysis of the eigentensor, giving the linear combinations of traits that describe the difference in the eigentensor. Lambda describes the amount of variation in the eigentensor that each eigenvector explains (with the proportion of variance in parentheses), with the trait loadings quantifying the contribution of each trait to differences in variance. Only the first eigentensor is presented with the full summary located in Table S7. Numbers in bold denote trait loadings higher than 0.25, suggesting they contributed to the difference in the eigenvector of the eigentensor. Plant architecture traits accounted for the greatest difference in genetic variance. Traits: 1 = Height, 2 = Stem length / width, 3 = # branches, 4 = Stem diameter, 5 = Leaf area, 6 = Perimeter^2^ / Area^2^, 7 = Circularity, 8 = # indents, 9 = Indent width, 10 = Indent depth.

| Eigen-tensor | $\alpha$ | Eigen-vector | $\lambda$ | Architecture traits | | | | | Leaf morphology traits | | | | |
| --- | --- | --- | --- | --- | --- | --- | --- | --- | --- | --- | --- | --- | --- |
|  |  |  |  | 1 | 2 | 3 | 4 | 5 | 6 | 7 | 8 | 9 | 10 |
| E1 | 1.23<br>(0.38) | $\mathbf{e}_{1,1}$ | -0.95<br>(0.61) | -0.24 | **-0.44** | 0.04 | **-0.50** | **-0.62** | 0.07 | -0.08 | 0.22 | -0.01 | -0.21 |
| | | $\mathbf{e}_{1,2}$ | -0.27<br>(0.17) | 0.24 | **-0.50** | **-0.82** | 0.07 | 0.12 | -0.02 | 0.02 | -0.06 | 0.04 | 0.01 |
| | | $\mathbf{e}_{1,3}$ | -0.08<br>(0.05) | **0.47** | -0.15 | 0.18 | **0.53** | **-0.58** | 0.05 | 0.14 | -0.01 | -0.22 | 0.21 |
| | | $\mathbf{e}_{1,4}$ | 0.08<br>(0.05) | 0.02 | -0.02 | -0.01 | **0.25** | 0.15 | **-0.41** | -0.10 | **0.72** | **-0.28** | **-0.37** |
| | | $\mathbf{e}_{1,5}$ | -0.06<br>(0.04) | 0.09 | **-0.70** | **0.52** | 0.03 | **0.46** | 0.12 | -0.04 | -0.03 | 0.00 | 0.05 |
| | | $\mathbf{e}_{1,6}$ | -0.04<br>(0.02) | **0.76** | 0.15 | 0.09 | **-0.59** | 0.08 | -0.17 | 0.04 | 0.08 | -0.02 | 0.02 |
| | | $\mathbf{e}_{1,7}$ | 0.03<br>(0.02) | -0.24 | -0.11 | 0.02 | -0.16 | 0.02 | **-0.56** | **0.50** | **-0.30** | **-0.47** | 0.17 |
| | | $\mathbf{e}_{1,8}$ | 0.02<br>(0.01) | 0.02 | -0.11 | 0.10 | 0.11 | -0.15 | **-0.68** | **-0.30** | -0.17 | **0.59** | 0.07 |
| | | $\mathbf{e}_{1,9}$ | 0.02<br>(0.01) | 0.14 | 0.01 | 0.07 | 0.11 | -0.03 | 0.00 | 0.09 | **-0.47** | -0.02 | **-0.86** |
| | | $\mathbf{e}_{1,10}$ | 0.00<br>(0.00) | 0.01 | 0.02 | -0.04 | -0.05 | -0.01 | -0.06 | **-0.78** | **-0.29** | **-0.54** | 0.08 |

The first eigentensor was dominated by the first eigenvector (***e***_1_,_1_), a linear combination of plant architectural traits and leaf area that explained 61% of the divergence in genetic variance between ecotypes for E1 (Table 2). The first three eigenvectors of E1 described divergence in genetic variance underlying different aspects of architecture traits and leaf size, accounting for 83% of the difference in genetic variance in E1. In contrast, differences in genetic variance underlying leaf shape traits (***e***_1_,_4_ and ***e***_1_,_7_ - ***e***_1_,_10_) together only explained 9% of the difference in genetic variance between ecotypes for E1 (Table 2). The coordinates of the ecotype **G** matrices in the space of the first eigentensor showed that divergence in genetic variance between the Headland ecotype and the Dune and Woodland ecotypes were responsible for the divergence in genetic variance described by the eigentensor (Figure 4A). Projection of ***e***_1_,_1_ through the original **G** matrices quantified the contribution of each ecotype to the major axis of divergence in genetic variance, with results providing further evidence that divergence in genetic variance was created by differences between the Headland ecotype and the Woodland and Dune ecotypes (Figure 4B). Therefore, the Headland ecotype has diverged the most from the other ecotypes in genetic variance for plant architecture and leaf size.

**Figure 4.**
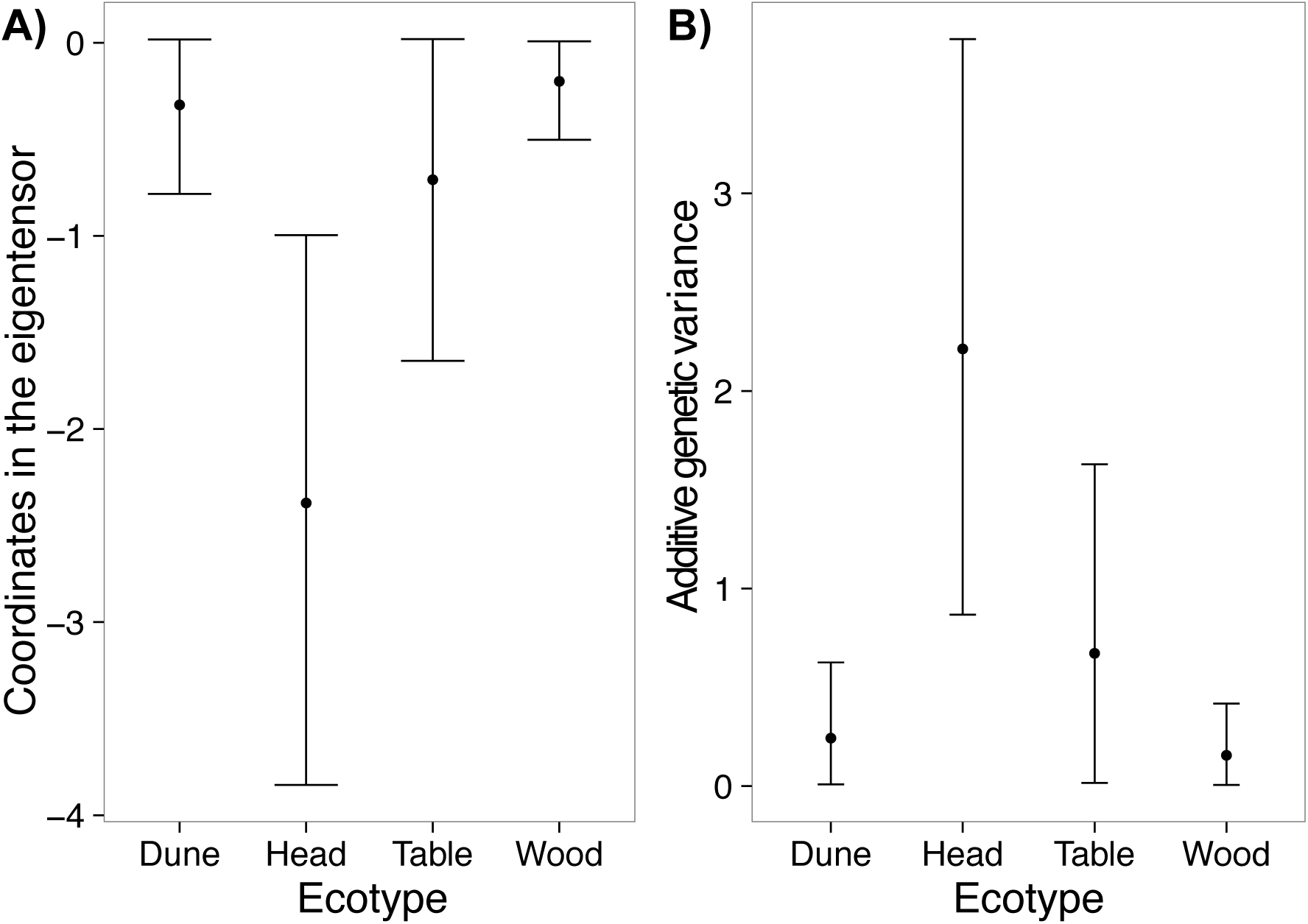
(**A**) Mean and 95% HPD intervals for the coordinates of each ecotype **G** matrix in the space of the first eigentensor (E1) of **G**. (**B**) Mean and 95% HPD intervals for the projection of the leading eigenvector from the first eigentensor of **G** through the original ecotype **G** matrices. The Headland ecotype showed strong divergence in additive genetic variance from the Dune and Woodland ecotypes.

#### Aligning divergence in phenotype mean with divergence in genetic variance

Projection of eigenvectors from the first eigentensor of **G** (***e***_1_,_1_ - ***e***_1_,_10_), through **D** quantified the alignment between divergence in genetic variance and divergence in phenotypic mean. The ***e***_1_,_1_ trait combination described greater divergence in mean than expected by chance (Figure 5), suggesting that substantial divergence in genetic variance was associated with divergence in phenotypic mean. However, two eigenvectors associated with leaf shape traits (***e***_1_,_4_ and ***e***_1_,_7_) also described a large amount of divergence in phenotypic mean, suggesting that small genetic changes were also associated with phenotypic divergence. These results suggest that evolution of continuous traits occurs when natural selection creates changes in the distribution of additive genetic variation.

**Figure 5.**
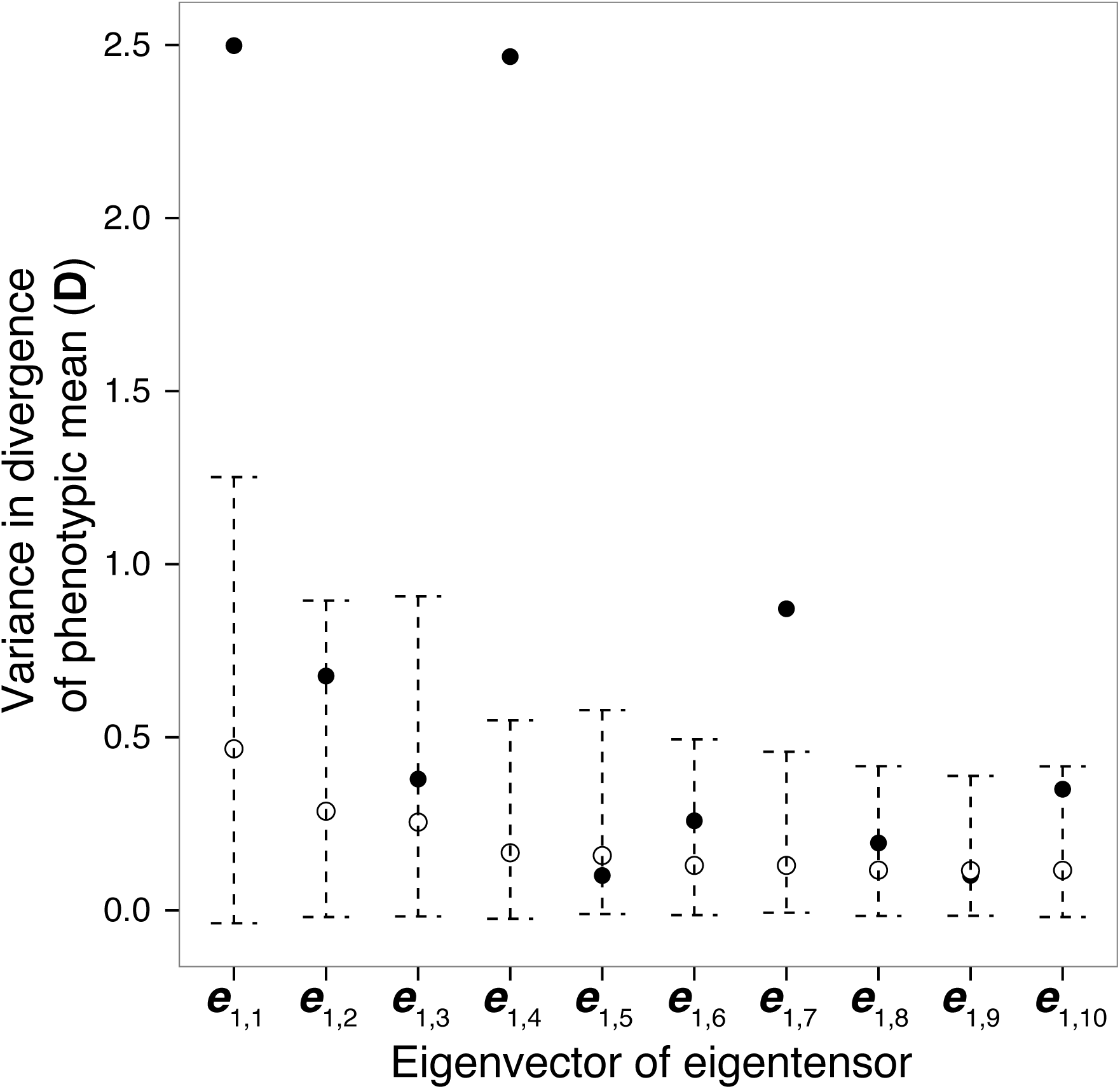
Projection of eigenvectors of the first eigentensor of **G** through the observed (filled circles) and randomised (unfilled circles) **D** matrices. Dashed lines represent 95% HPD intervals for the projections through the randomised **D** matrices. Eigenvector one showed a strong association with divergence in mean phenotype. However, eigenvectors four and seven also explained a large amount of variance in divergence, suggesting small changes in genetic variance also explained large variance in divergence.

Adaptation is expected to move along the line of greatest genetic variance (***g***_max_) (29), but ***g***_max_ itself might change during the early stages of adaptive divergence if natural selection or genetic drift change allele frequencies in genes responsible for trait variation in the system. Eigentensors represent differences in the entire space of genetic variance, so to explore whether changes in specific axes of genetic variance aligned with divergence in phenotype mean we quantified the alignment between changes in the orientation of ecotypic ***g***_max_ and ***g***_2_, with divergence in phenotype mean. First we converted ***g***_max_ and ***g***_2_ into (co)variance matrices using

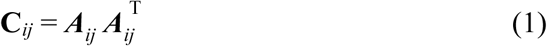

where *A_ij_* represents the *i*th vector of **G** (***g***_max_ or ***g***_2_) for the *j*th ecotype. This approach resulted in four (co)variance matrices that represented ***g***_max_ for the four ecotypes (**C**_***g***max_), and four matrices that represent ***g***_2_ (**C**_***g***2_). Conducting separate covariance tensor analyses on **C**_***g***max_ and **C**_***g***2_ gave the divergence in the orientation of ***g***_max_ and ***g***_2_ between ecotypes. The first eigentensor of both tensor analyses captured large differences between ecotypes for both eigenvectors of G, where E1 described 51% of divergence in ***g***_max_ and 48% of divergence in ***g***_2_. The corresponding eigenvectors of eigentensors represented the axes of greatest divergence in the orientation of ***g***_max_ and ***g***_2_. The first two eigenvectors of E1 (*e*_1_,_1_ and *e*_1_,_2_) for ***g***_max_ and ***g***_2_ described most of the difference in variance explained by the first eigentensor (***g***_max_ e_1_,_1_ = 50% and ***e***_1_,_2_ = 42%; g_2_ ***e***_1_,_1_ = 50% and ***e***_1_,_2_ = 38%).

Projection of the eigenvectors from the first eigentensor of **C**_***g***max_ and **C**_***g***2_, through the observed and random **D** matrices (using equation 6) quantified the amount of divergence in mean phenotype associated with divergence in the orientation of the major eigenvectors of ecotype **G**. For ***g***_max_, both ***e***_1_,_1_ and ***e***_1_,_2_ accounted for more divergence in phenotype mean than expected by chance (Figure 6A). For ***g***_2_, only eigenvectors associated with very small divergence in orientation (***e***_1_,_3_, ***e***_1_,_4_ and ***e***_1_,_6_) described a similar amount of divergence in phenotype mean (Figure 6C). Therefore, strong differences in the orientation of ***g***_max_, but not ***g***_2_ aligned with divergence in mean phenotype, suggesting that additive genetic variance has aligned with the direction of natural selection for each ecotype.

**Figure 6.**
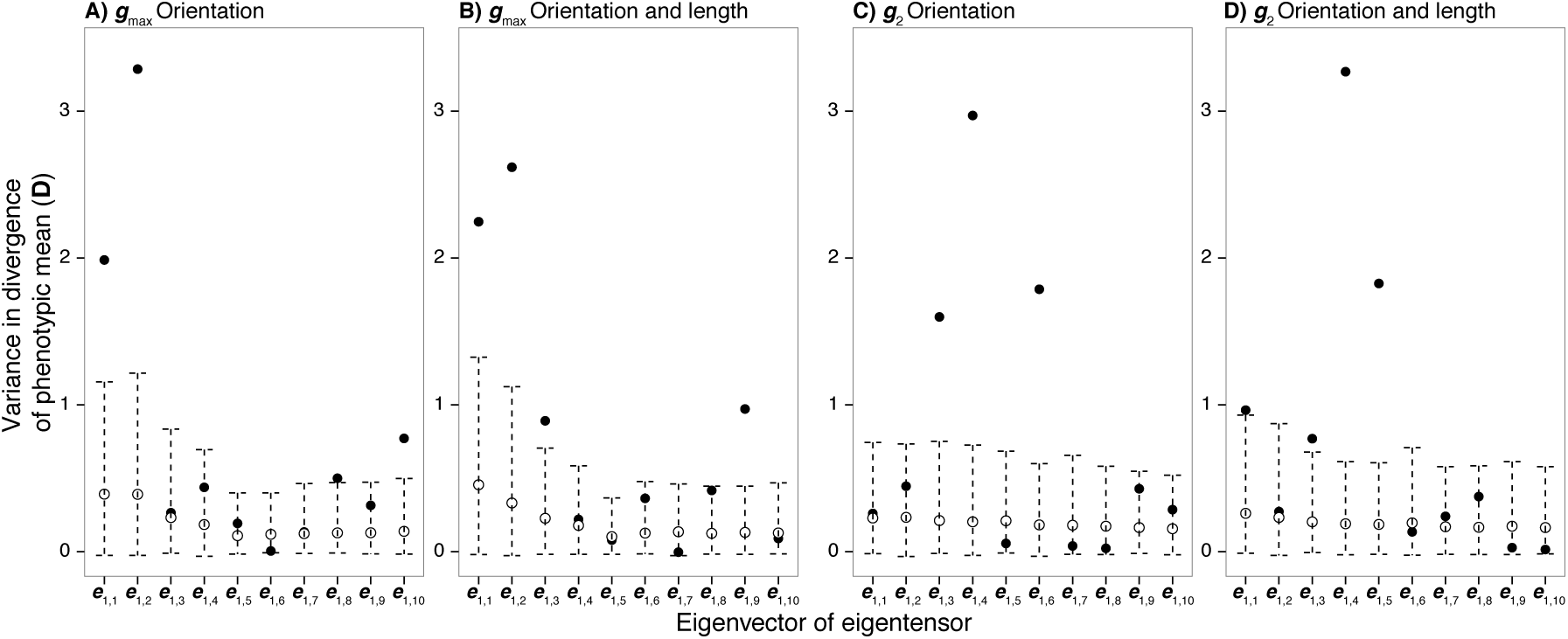
Projection of eigenvectors from the first eigentensor taken from tensor analyses conducted on (**A**) **C**_***g***__max_, (**B**) **C**′_***g***max_, (**C**) **C**_***g***2_ and (**D**) **C**′_***g***2_, through the observed (filled circles) and random (unfilled circles) **D** matrices. Confidence intervals represent 95% HPD intervals for the projection through the random **D** matrices. Divergence in the orientation of ***g***_max_, but not ***g***_2_ described strong divergence in mean phenotype. Including divergence in the length of ***g***_max_ and ***g***_2_ did not change the result, suggesting that differences in the amount of genetic variance in each eigenvector of **G** was not associated with phenotypic divergence.

Differences between ecotypes in eigenvectors of **G** can also be due to differences in length. To identify whether differences in the length of the original eigenvectors contributed to describing divergence in phenotype mean we repeated the analysis for equation 1 and included the length of the original eigenvector using

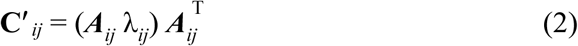

where *A_ij_* represents the *i*th vector of **G** (***g***_max_ or ***g***_2_) for the *j*th ecotype, with λ*_ij_*. the associated eigenvalue. **C**′_*ij*_ then represents differences in the orientation and length of ***g***_max_ and _***g***2_ (**C**′ *_g_*_max_ and **C**′ *_g_*_2_). A tensor analysis on **C**′ *_g_*_max_ and **C**′ *_g_*_2_ found that E1 described 85% and 65% of the variation in the orientation and length of ***g***_max_ and ***g***_2_, respectively. The first two eigenvectors described most of the difference in E1 for both ***g***_max_ (*e_1,2_* = 84% and ***e***_1_,_2_ = 12%) and ***g***_2_ (***e***_1,2_ = 50% and ***e***_1,2_ = 49%). Projection of ***e***_1_,_1_ and ***e***_1_,_2_ for **C′**_***g***max_ and **C′**_***g***2_, through the observed and random *D* matrices showed that including information about divergence in the length of the eigenvectors of G did not change the results appreciably. For **C′** *_***g***_*_max_ (Figure 6B), ***e***_1_,_1_ and ***e***_1_,_2_ together described less divergence in mean, whereas eigenvectors that described only very minor changes in the length and orientation of ***g***_max_ described more divergence in mean than for orientation alone (Figure 6A). Therefore, differences in orientation, but not the length of ***g***_max_ described divergence in mean phenotype. For **C′** _***g***2_ (Figure 6D), ***e***_1_,_1_ and e_1_,_2_ together described more divergence in mean phenotype than orientation alone (Figure 6C), suggesting that some differences in the length, but not orientation of ***g***_2_ was associated with divergence in mean phenotype.

## Discussion

The conceptual framework underlying Simpson’s adaptive landscape connects evolutionary processes such as natural selection and genetic drift, to species diversification during adaptive radiation (1, 51, 52). The conceptual adaptive landscape is represented as a bivariate phenotypic space overlaid with a fitness surface where topographical peaks represent fitness optima. Adaptation on the adaptive landscape occurs when a population climbs a fitness peak in phenotypic space, with divergence occurring when populations climb different peaks (52). Movement across an adaptive landscape can occur when selection on additive genetic variance shifts the population mean towards a fitness peak (1). If a common ancestor colonizes multiple environments, the adaptive landscape will be represented by the same phenotypic space with a different fitness surface for each environment. Natural selection in each environment will pull the phenotype mean towards fitness peaks in different areas of phenotypic space, creating adaptive divergence. However, when the axis of greatest additive genetic variance aligns away from the direction of selection, adaptation is expected to take an indirect route towards the fitness peak, constraining the rate of adaptation (28, 29).

Incorporating the concept of the adaptive landscape into an empirical context has a number of challenges. Few empirical studies have used the adaptive landscape to investigate patterns and processes underlying diversification (but see 53, 54), and incorporating the role of the underlying genetic variance in patterns of divergence are also rare (Chenoweth et al 2010). Furthermore, adaptive radiations typically involve suites of traits, and characterizing divergence in phenotypic mean and genetic variance is more complex than the conceptual representations of two phenotypic traits and fitness. Here, we have shown how to use consistent ecotypic divergence among replicate populations of *S. lautus* to investigate adaptive divergence across the adaptive landscape by exploring the association between divergence in multivariate phenotype mean and divergence in genetic variance in an early adaptive radiation.

### Multivariate divergence in mean phenotype and genetic variance among ecotypes

Multivariate phenotypic divergence was stronger between ecotypes than within ecotypes, indicating environment specific phenotypes have arisen due to similar natural selection regimes on populations that share similar habitats (55, 56). Phenotypic divergence was very strong in two axes, separating Tableland and Woodland from the remaining ecotypes in one axis, and Dune and Headland from the remaining ecotypes in the other axis (Figure 1). Additional evidence for natural selection creating divergence comes from the repeated and independent evolution of parapatric pairs of Dune and Headland populations along the coastline, with pairs separated by large distances (tens to hundreds of kilometres) (46, 47, 57). These patterns of replicated evolution across a wide geographical scale are similar to those documented in sticklebacks (14), African cichlids (48) and *Anolis* lizards (16).

We also found divergence in the distribution of additive genetic variance underlying the same morphological traits, driven largely by the Headland ecotype. Heritabilities were very different between ecotypes, with higher values observed for architecture than leaf traits, especially in the Headland ecotype. Divergence in multivariate genetic variance was captured by one major axis representing plant architecture traits, while leaf morphology traits explained very little divergence in genetic variance. Ecotypes therefore varied both in the magnitude of genetic variance in the traits included in our study, and in distribution of genetic variance across multivariate phenotypic space. In particular, the Headland ecotype had more genetic variance that was highly concentrated into a few trait combinations.

The Tableland and Headland ecotypes showed higher heritabilities and higher genetic variance in linear combinations of architecture traits, which was reflected by contrasting architectural phenotypes. The prostrate form of the Headland ecotype is especially contrasting to the other ecotypes and is typical of many plants occupying exposed environments subject to strong winds (10, 58). Highly diverged genetic variance underlying plant architecture in the Headland ecotype suggests that differences in natural selection required strong genetic changes to create the prostrate phenotype beneficial in the headland environment. Plant architecture for the Tableland ecotype was similar to the Dune and Woodland ecotypes, but more extreme in terms of overall plant size, potentially due to selection for larger plant size in a rainforest environment (59–61). Therefore, natural selection along genetic pathways common to the Dune, Woodland and Tableland ecotypes may have created larger plants in the Tableland environment.

### Aligning divergence in phenotype mean with divergence in genetic variance

During the initial stages of adaptive divergence, the association between the direction of natural selection on multivariate phenotypes and the distribution of genetic variance is expected to determine evolutionary trajectories. However, over time the direction of selection alone is expected to dominate adaptive divergence between habitats, regardless of the underlying genetic architecture (25, 29). In our experiment, the presence of strong divergence in **G** itself argues against making an implicit assumption of a stable ancestral **G** matrix and its influence on divergence in phenotype. Our results are consistent with several studies that found differences in **G** following very recent divergence (62–64), suggesting that rapid adaptive divergence may occur when new environments are colonized and the distribution of genetic variance aligns rapidly with the direction of natural selection.

For recently derived ecotypes of *S. lautus*, we have shown that divergence in **G** aligned with divergence in phenotypic mean, suggesting that during adaptation natural selection has changed the distribution of genetic variance. More specifically, changes in the orientation of ***g***_max_ but not the orientation of ***g***_2_ were associated with changes in phenotype mean. Therefore, the linear combination of traits with the greatest genetic variance may have changed orientation towards the direction of selection, facilitating rapid adaptive divergence. Perhaps the simplest explanation for finding an association between divergence in mean and divergence in the major axis of genetic variance would involve a selection response based on alleles segregating at low frequency. Initially rare alleles maintained by mutation-selection balance would contribute little to genetic variance (36, 37). After colonization of a novel habitat some may become beneficial and rise in frequency during adaptation, contributing more substantially to levels of additive genetic variation in traits under selection (65).

Trait combinations with strong contributions from plant architecture traits accounted for most of the divergence in genetic variance. There is some evidence to suggest that plant architecture traits may be controlled by many genes of small effect (66, 67), and in mammals body size is determined by numerous genes of small effect, many of which are at low frequency (Kemper et al. 2012). Divergence among these ecotypes in plant architecture might have been based to some extent on such alleles. In contrast, linear combinations of leaf morphology traits described much smaller fractions of the total divergence in genetic variance. Leaves are responsible for the water-energy balance of plants, which is likely to require specialized traits or combinations of traits within each environment (59). Previous research has found a relatively small number of loci control leaf morphology in *Arabidopsis* and *Populus* (68, 69). If selection on a small number of alleles controlling leaf shape is strong, then alleles will move rapidly towards higher frequency, reducing the genetic variance within each ecotype for leaf traits underlying adaptive divergence.

### Conclusions

If genetic correlations between traits bias evolution then it is difficult to see how rapid adaptive divergence leads to adaptive radiation. Our results suggest that alleles present in standing genetic variation (possibly rare) may become beneficial when new environments are colonized, increasing their frequency in the population and aligning the distribution of genetic variation with the direction of natural selection. Adaptation from standing genetic variation can be rapid (70), suggesting that natural selection shaping the distribution of genetic variation can provide the mechanism for understanding how adaptive radiation proceeds. Ascension of adaptive peaks on the adaptive landscape can then occur when different alleles are favoured in different environments, aligning the direction of greatest genetic variation with the direction of selection in each environment. The presence of repeated adaptation to similar environments further suggests that natural selection has favoured the same alleles in similar environments (46), driving the adaptive radiation of ecotypes into multiple contrasting environments.

## Methods

### Growth protocol and phenotype measurement

To grow seeds in the glasshouse we induced seed germination by scarifying each seed with a razor blade, placing them on moist filter paper in glass petri dishes and leaving them in the dark for 2 days. Seedlings were then transferred to a controlled-temperature room at 25°C on a 50:50 day:night light cycle. After one week, seedlings were taken to the glasshouse and planted into 137mm round pots in glasshouse experiment one and 85mm square pots in glasshouse experiment two. Pots contained soil (70% pine bark: 30% coco peat) with 5kg/m^3^ osmocote slow-release fertiliser and 830g/m^3^ Suscon Maxi soil insecticide. We performed controlled crosses between plants by rubbing two mature flower heads together over successive days, allowing each flower to receive and donate pollen. Seeds were collected once mature and stored at 4°C until required for subsequent experiments.

Ten morphological measurements were taken using identical methods in both glasshouse experiments. Architectural measurements were recorded to the nearest mm with a ruler and included plant vegetative height, plant width at the widest point, plant width at the narrowest point and main stem length. Secondary branches were counted and main stem diameter was measured with callipers one inch from ground level. We divided the main stem length by an average of the two width measurements to get an architectural measurement that encompassed plant growth habit; low values signified a prostrate plant and high values indicated a tall, erect plant. One young but fully expanded leaf was taken from each plant and scanned using a flatbed scanner. Morphometric data of all scanned leaves was extracted using the program Lamina (71). Traits produced by Lamina included leaf area, perimeter, circularity, indent (serrations and lobes) width, indent depth and indent number. Perimeter squared divided by area squared was calculated as a measure of leaf complexity. Indent number was divided by perimeter to calculate indent density along the margin of the leaf.

Phenotypic traits for both experiments were normally distributed and because traits were measured on different scales we standardized traits within each ecotype to a mean of zero and a standard deviation of one. Scaling prevented traits dominating the eigenstructure of our analyses due to differences in measurement units (72, 73).

### Divergence in multivariate mean among ecotypes

To investigate divergence in mean phenotype we conducted a MANOVA using the nested linear model,

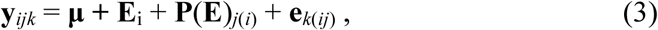

where **μ** was the intercept and the sources of variation in the experiment were represented by ecotype (**E***_i_*), populations nested within ecotype (**P**(**E**)_*j*(*i*)_) and the residual error (**e***_k_*(*_ij_*)). The ten phenotypic traits were fitted as a multivariate response variable (**y***_ijk_*). From the MANOVA we calculated the **D** matrix by extracting the sums of squares and cross-product matrices for ecotype (SSCP_H_) and population (SSCP_E_) and calculated the mean squares matrices by standardising by their associated degrees of freedom (MS_H_ = SSCP_H_ / 3; MS_E_ = SSCP_E_ / 12). We then calculated **D** using

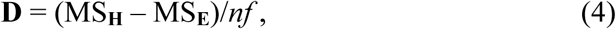

where *nf* represented the number of individuals sampled from each ecotype in a balanced design. Given slight differences in numbers between ecotypes we calculated *nf* using equation 9 in Martin et al. (2008). We used this method for calculating **D** to isolate the divergence between ecotypes from differences between populations and use the resultant (co)variance matrix to compare with differences in genetic variance (74). Eigenanalysis of **D** gave the linear combination of traits that explained divergence in mean phenotype between ecotypes with the associated eigenvalue representing the amount of divergence. Three degrees of freedom at the ecotype and population levels gave a maximum of three non-zero eigenvectors. Calculating the scores of **D** for each population using the first two vectors of divergence visualised separation between populations in the major axes of phenotypic divergence. To compare **D** to a null distribution we created a null expectation for **D** by randomising individual phenotypes between ecotypes 1,000 times and re-calculating **D** for each randomisation. Divergence in genetic variance could then be compared to both the observed and randomised **D** matrices.

### Estimation of multivariate genetic variance components

We set out to estimate genetic variance for each population, but due to modest replication of phenotyped offspring we pooled the data for the populations for each ecotype and estimated genetic variance for each ecotype separately. To estimate the genetic variance components for each ecotype separately we used the R package ‘MCMCglmm’ (75) to fit a sire model,

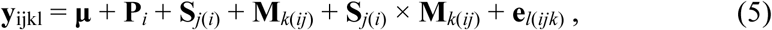

where replicate population (**P***_i_*), and the intercept (**μ**) were fitted as fixed effects. Including replicate population as a fixed effect removed the variance due to difference in means between populations (i.e., population divergence), within an ecotype. Sire (**S***_j_*_(*i*)_), dam (**M***_K_*_(*ij*)_), and the sire × dam interaction (**S***_j(i)_* × **M***_K_*_(*ij*)_) were fitted as random effects and ***e***_*l*_(*_ijk_*) was the residual variance. Phenotype measurements for each ecotype (**y***_ijkl_*) were standardized to a mean of zero and standard deviation of one before being entered as a multivariate response. Each model was run for 2,100,000 Marcov chain Monte Carlo (MCMC) sampling iterations, which included a burn-in of 100,000 MCMC iterations and a thinning interval of 2,000 MCMC iterations. We used a Cauchy prior distribution (76) and to examine the sensitivity of the prior we adjusted the parameters to excessively large and small values, making sure the model output remained stable. All models converged with autocorrelation below 0.05 between MCMC sampling iterations and the effective sample size exceeded 85% of the total number of samples for all parameters estimated. We then calculated the additive genetic variance matrix, **G** (**G**_obs_) as four times the sire variance (**S***_j_*_(*i*)_).

Bayesian estimation of genetic variance components has the benefit of being able to carry uncertainty throughout the analysis by applying the same transformation to each MCMC sampling iteration (75, 77). However, variance component estimates are constrained to be greater than zero (positive-definite) and therefore, testing for significant estimates of genetic variance requires comparing **G**_obs_ to a suitable null expectation. We used random **G** matrices (**G**_ran_) as the null expectation, which we created by randomising phenotype with respect to parentage (within ecotypes) and re-running model 5. We conducted 1,000 randomisations within each ecotype and ran a separate model for each randomised dataset. Due to the significant computer running time required we kept the thinning interval and burn-in period identical, but reduced the total number of iterations such that we sampled the smallest number of iterations that would give us a reliable estimate of the posterior mean. The minimum total number of iterations was calculated from our observed models of **G** by estimating the mean for a variance component from an increasing number of iterations until the accuracy of our estimate of the mean reached an asymptote. We checked convergence for the randomised **G** models with all models showing no autocorrelation between MCMC samples and effective sample sizes greater than 85% of the number of iterations saved. Taking the posterior mean **G** for each random model gave us 1,000 randomized G matrices. We then compared the variance explained by **G**_obs_ to **G**_ran_ for several analyses outlined below.

### Characterising ***G***

To characterise **G**, we estimated the univariate heritability for each trait. Traits were standardised to a variance of one within each ecotype separately and therefore, the diagonal elements of our observed **G** represented heritabilities. To identify whether our estimated heritabilities were higher than expected by sampling error, we compared the heritabilities taken from our posterior mean observed **G** with those taken from our randomized **G** matrices. Taking the diagonal elements for each of our 1,000 randomised **G** matrices gave 1,000 random estimates of the univariate heritabilities. If the heritability estimates from our observed **G** were higher than the 95% Highest Posterior Density (HPD) interval for the randomized heritability distribution, we took this as evidence that our observed heritabilities were higher than expected by sampling error.

To explore the distribution of genetic variance we used eigenanalyses on the posterior mean of our observed **G**. The distribution of genetic variance among eigenvectors describes the shape of multivariate genetic variance, where fewer eigenvectors with relatively high genetic variance denote a more elliptical **G**. The linear combination of traits with the most genetic variance then describes the direction of greatest multivariate genetic variance. To quantify whether eigenvectors explained more variance than expected by sampling error, we compared eigenanalyses conducted on **G**_obs_ with **G**_ran_. We conducted an eigenanalysis on each of the 1,000 random **G** matrices and saved the eigenvalues, which gave the distribution of the null expectation for each eigenvector of **G**. If the eigenvalues from our observed **G** were higher the upper 95% confidence interval for the distribution of the randomised eigenvalues, then our observed eigenvectors explained more variance than expected due to sampling error.

### Divergence in multivariate genetic variance among ecotypes

To investigate divergence in genetic (co)variance matrices between ecotypes we used the covariance tensor approach, which quantifies differences between multiple matrices and uses eigenanalysis to identify how traits contribute to these differences. The tensor analysis is a three-step process; first we constructed the **S** matrix, which contained the variances and covariances between all matrix elements (summarised in 77, 78). Second, eigenanalysis of **S** gave the eigentensors, which are matrices that explained the most difference in variance between ecotype matrices, with their associated eigenvalue quantifying the amount of variance in the difference attributed to each eigentensor. Third, a second eigenanalysis, conducted on the eigentensors, gives the eigenvectors that describe the distribution of the variation within each eigentensor, corresponding to the linear combination of traits that explain the difference between the original matrices (77, 78).

To determine the eigentensors associated with significant differences in genetic variation, we compared the difference in genetic variance explained by tensor analyses on our observed and randomised **G** matrices (77). For the random tensor analysis we conducted a covariance tensor on each randomised model of **G** and saved the **S** matrix, giving 1,000 random estimates of **S**. We then projected the eigenvectors of **S** from the tensor analysis of **G**_obs_, through the random S matrices, quantifying the amount of difference in variance in the randomised **G** matrices explained by each of the observed eigenvectors. We then compared the distribution of random eigenvalues of **S** to the observed eigenvalues of **S**. Where the observed eigenvalues of **S** exceeded the 95% HPD intervals of the random distribution was interpreted as an eigentensor that accounted for significant differences in variation between ecotype matrices.

To investigate the contribution of each ecotype to differences in variance described by the eigentensors we used two methods. First, we calculated the frobenius product between the original ecotype **G** matrices and the eigentensor, which gave the coordinates of each ecotype in the space of the significant eigentensors. Where ecotypes show differences in the coordinates reveals which ecotypes contribute to the differences in the eigentensor. Second, matrix projection of the leading eigenvectors of the eigentensors, through the original matrices, examines how much of the difference in variance explained by the leading eigenvectors is attributable to each ecotype.

### Aligning divergence in phenotype mean with divergence in genetic variance

To investigate whether divergence in variance was associated with divergence in mean multivariate phenotype we compared the eigentensor of **G** with **D**. To do so, we projected the eigenvectors of eigentensors through **D** using

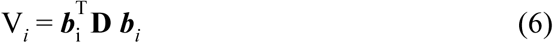

where is the ***b****_i_ i*th normalised eigenvector of the first eigentensor and **D** is the divergence matrix. V*_i_* is then the amount of divergence in mean phenotype associated with each eigenvector of the eigentensor. We used equation 6 on the 1,000 MCMC iterations of our observed **G**, and the 1,000 randomisations of **D** to carry through the uncertainty for both **D** and the eigentensor of **G**. First, we took the first eigentensor from tensor analyses conducted on each MCMC iteration of **G**, giving 1,000 estimates of the eigentensor. We then projected the eigenvectors of these eigentensors through the 1,000 randomisations of **D**. If V*_i_* explained more divergence than expected by chance, the observed projection would fall outside the 95% confidence intervals of the null distribution.

## Acknowledgments

We would like to thank Carol Palmer, who was integral in the collection of the phenotype data. We would also like to thank the University of Queensland glasshouse staff for their help and support. We are grateful to Tom Richards for his help with field sampling and providing comments on previous manuscripts. We would also like to thank Federico Roda, Gail Walter, Kristylee Marr and James Donohoe for their help with field sampling. Scott Walter provided important help in the glasshouse. Emma Hine provided insightful discussions during analysis. This project was funded by Australian Research Council grant DP0986172. Seeds were collected in NSW national parks under permit number S12084.

## Statement of authorship

GW, JDA, MB and DO designed the research; GW conducted field sampling and glasshouse work; GW and JDA analysed data with input from DO and MB; GW, JDA, MB and DO wrote the manuscript. All authors declare no conflict of interest.

